# Induction of Bacteriophage PinR Facilitates the Evolution of Antibiotic Resistance

**DOI:** 10.1101/2022.03.29.486199

**Authors:** Le Zhang, Yunpeng Guan, Qian Peter Su, Yan Liao, Iain Duggin, Dayong Jin

**Affiliations:** Institute for biomedical materials and devices, University of Technology Sydney, Ultimo, Australia; Department of biomedical engineering, Southern University of Science and Technology, Shenzhen, China; Australian Institute for Microbiology & Infection, University of Technology Sydney, Ultimo, Australia

## Abstract

A recent work reports that the single treatment of β-lactams can cause a SOS-independent superfast evolution of multi-drug resistance in the DNA repair deficiency *Escherichia coli* (*E. coli*), but the mechanism is not yet clear. Here, we find that the induction of PinR, a lambdoid prophage Rac, is involved in this process and facilitates the evolution of antibiotic resistance in DNA repair deficiency bacteria through the repression on the transcription of antioxidative genes and the thereafter ROS burst in cells. It is highlighted that the bacteriophage PinR can orchestrate the mutagenesis induced by the overaccumulation of ROS in cells. More importantly, we for the first time demonstrate that the deletion of *pinR* can avoid the rapid evolution of antibiotic resistance induced by either the single or long-term exposure to antibiotics, while strategies to target RecA, *e*.*g*., the inactivation on RecA, can be safely implemented to disarm the bacterial resistance to other antibiotics. Therefore, from a drug development perspective, our work suggests future studies on the “evolutionary potentiators” towards a safe and more effective strategy to be developed for infectious disease treatment.

## Main

Antibiotics can kill bacteria by inducing the DNA damage and repressing the DNA repair in cells (1), but bacteria can repair the DNA damage via a series of intrinsic pathways including the SOS response that supports cell survival upon DNA damage (2-5). The master regulator of the SOS response is RecA, which can enable bacteria to repair DNA damage and help bacteria to drive the development and spreading of antibiotic resistance determinants (6-8). Thus, it has been believed that deactivating the RecA may disarm the bacterial resistance to antibiotics. However, as a double-edged sword, the deficiency of DNA repair, such as a result of RecA inactivation, may also increase the drug resistance-related mutagenesis induced by the exposure to antibiotics (9,10). Notably, our recent work reports that a single treatment of β-lactams can cause a SOS-independent superfast evolution of multi-drug resistance in the DNA repair deficiency *E. coli* MG1655 (*ΔrecA* strain) (11), but the mechanism is not yet clear.

To better understand the mechanism, we here treated the wild type and the *ΔrecA* strain with a single exposure to ampicillin at 50 µg/mL for 8 hours. In line with our previous findings, a superfast evolution of antibiotic resistance was determined in the *ΔrecA* strain (Fig. S1A and B) (11). We explored the transcriptomic changes in surviving isolates induced by ampicillin, and found that the single treatment of ampicillin markedly affected the transcriptomic profile of the wild type and the *ΔrecA* strain compared to that of untreated control (Fig. 1A), with changes to the expression of 161 and 248 coding sequences (log_2_FC > 2 and *P* value < 0.05), respectively. However, the principal component analysis (PCA) showed that the effect of ampicillin on the *ΔrecA* strain remarkably differed to that of the wild type strain (Fig. 1B). Moreover, Venn diagrams confirmed that there were 115 and 202 genes specifically regulated by the exposure to ampicillin in the wild type and the *ΔrecA* strain, respectively (Fig. 1C). Because the formation of tolerance was observed in the wild type strain (Fig. S1C), these results collectively indicated that the drug exposure-induced differential regulation of bacterial transcriptomes led to different evolutionary directions.

**Figure 1.**
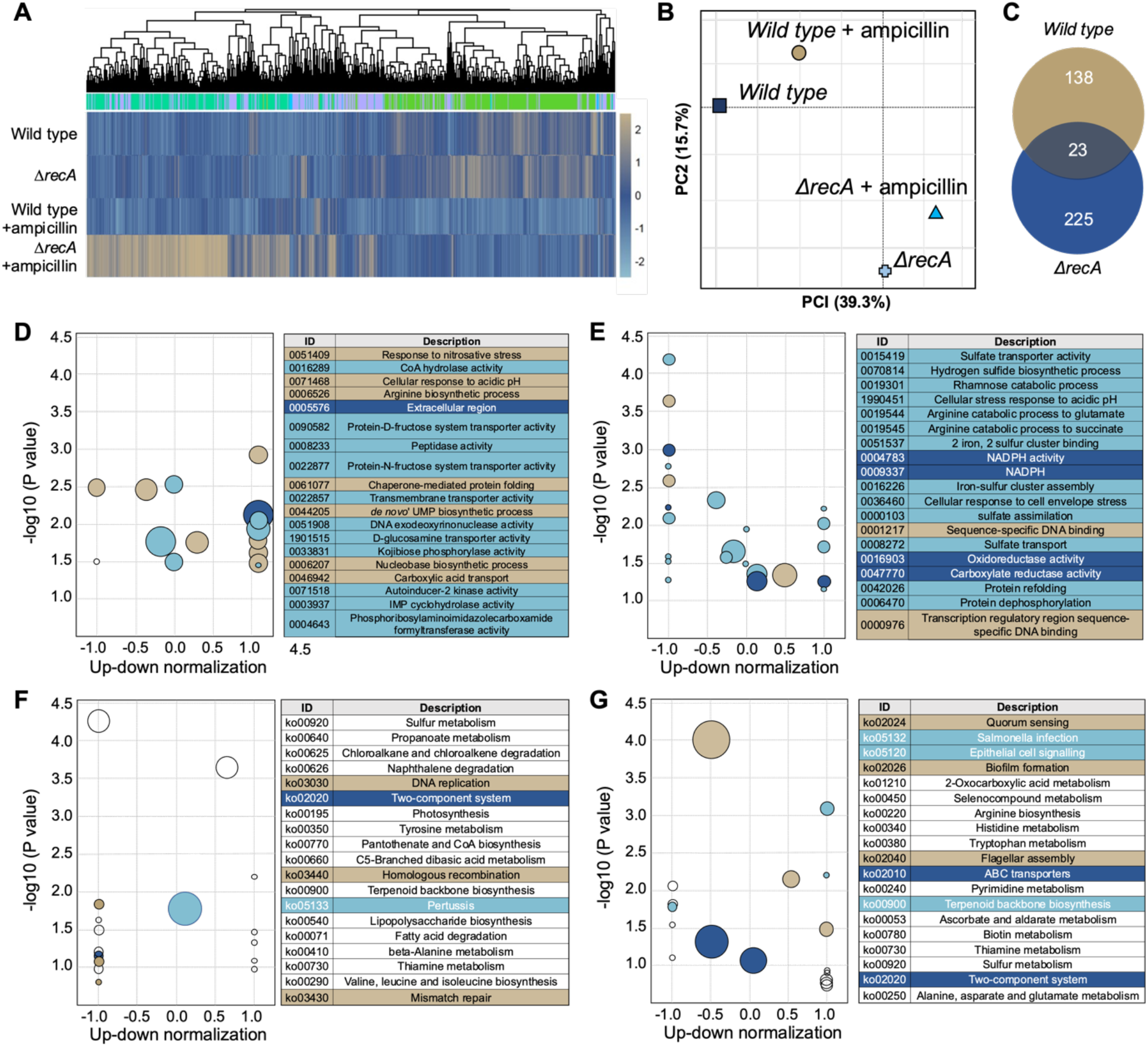
Transcriptional response of the wild type and the *ΔrecA* strain after a single treatment with ampicillin at 50 μg/ml for 8 hours. (A) The clustered heatmap of relative expression of coding sequences in the wild type and the *ΔrecA* strain with large fold changes and statistical significance (log_2_FC > 2 and *P* value < 0.05). (B) Bi-plot of the principal-component analysis (PCA) of normalized read counts for all strains. (C) Venn diagrams of differentially expressed genes (log_2_FC > 2). GO analysis scatter plot of different genes in the wild type (D) and the *ΔrecA* strain (E). KEGG pathway enrichment scatter plot of different genes in the wild type (F) and the *ΔrecA* strain (G). Top 20 enrichment pathways are listed in the GO and KEGG enrichment analysis. The global transcriptome sequencing was performed with two repeats in each group.

To study the different evolutionary trajectory, genes that were specifically regulated by ampicillin in the wild type and the *ΔrecA* strain were characterized. Genome-wide expression changes are visualized as volcano plots (Fig. S2) to identify specific genes with large fold changes and statistical significance (log_2_FC > 2 and *P* value < 0.05). Differential expression of genes related to certain biological functions defined by Gene Ontology (GO) enrichment analysis is shown in Fig. 1D and E. Kyoto Encyclopaedia of Genes and Genomes (KEGG) pathway analysis is shown in Fig. 1F and G. Overall, ampicillin had a major effect on the tolerance-related pathways in the wild type strain, including the quorum sensing, flagellar assembly, biofilm formation, and bacterial chemotaxis (12-14). In comparison, two functional categories were uniquely regulated in the *ΔrecA* strain, including the oxidative stress response, such as the sulfate transporter activity, iron-sulfur cluster assembly, oxidoreductase activity and carboxylate reductase activity, and the DNA damage response, such as the cellular response to DNA damage stimulus, DNA repair and recombinase activity.

Functional groups corresponding to biological processes were manually curated and visualized. First, the single treatment with ampicillin caused a significant downregulation of antioxidative genes transcription in the *ΔrecA* strain, including *cysJ, cysI, cysH, soda* and *sufD* (Fig. 2A), which indicated an overaccumulation of reactive oxygen species (ROS) in cells. The evolution of antibiotic resistance caused by the *in vivo* and *in vitro* elevated oxidative stress has been widely reported, since the induction of mutagenesis can be stimulated by the overproduction of ROS throughout the administration of antibiotics (15). We therefore asked whether the overproduction of ROS resulted in the superfast evolution of antibiotic resistance in the *ΔrecA* strain. To test this hypothesis, we added 50 mM glutathione (GSH), a natural antioxidative compound, into the culture medium and treated them with ampicillin at 50μg/ml for 8 hours. It is of significance that the addition of GSH prevented the evolution of resistance to ampicillin in the *ΔrecA* strain (Fig. 2B), and more importantly, it did not impair the bactericidal efficacy of ampicillin (Fig. 2C). We sequenced the surviving isolates and further found that the addition of GSH inhibited drug resistance-related DNA mutations in the *ΔrecA* strain, which could be detected in the *ΔrecA* resistant isolates including the gene *ampC* and *acrB* (Fig. 2D and Table S1). Taken together, these findings showed that the ROS burst is supposed to be a driver of the evolution of antibiotic resistance in DNA repair deficiency bacteria, but it is not the mechanism of death in ampicillin-treated cells.

**Figure 2.**
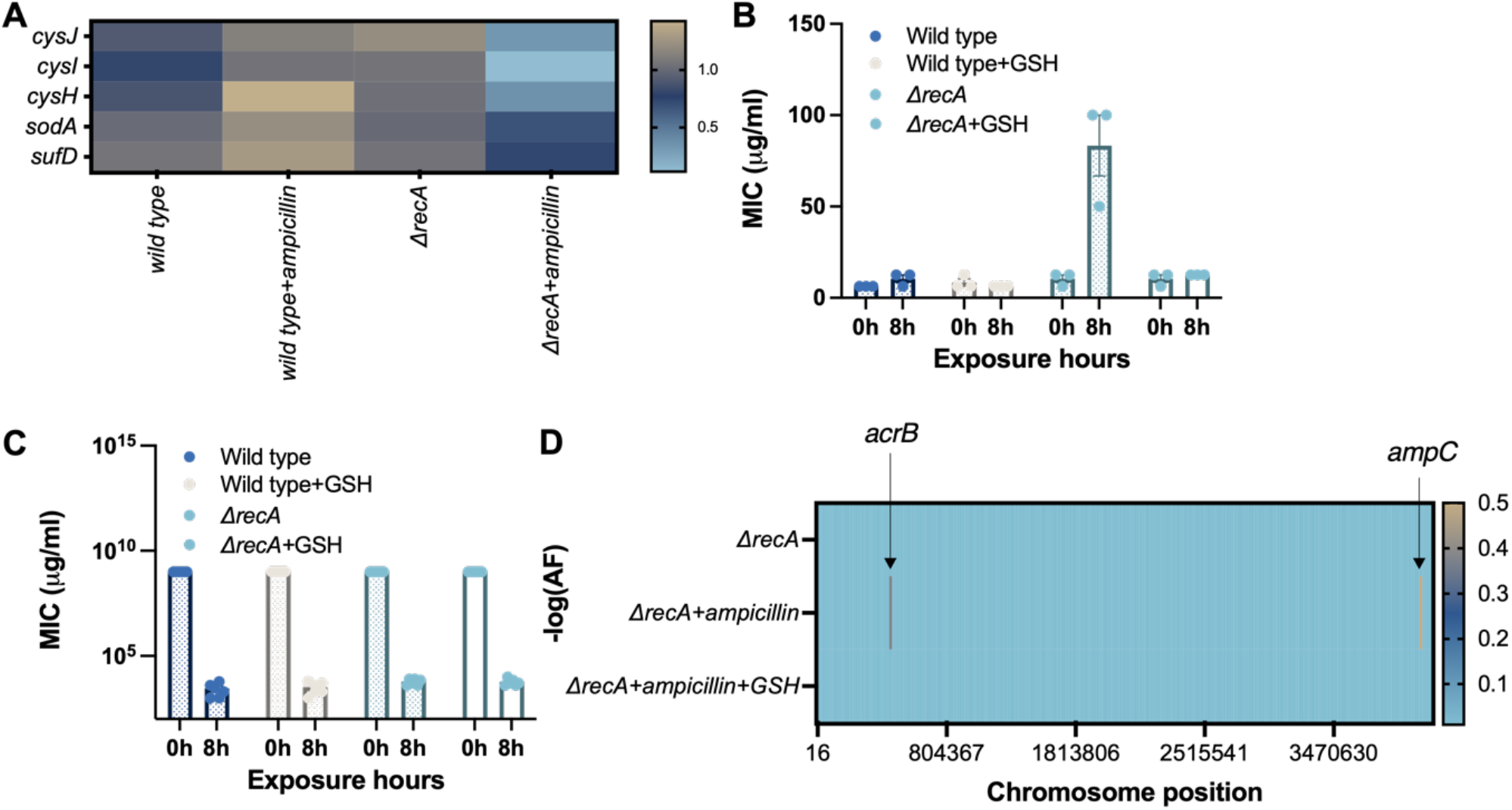
Overaccumulation of ROS is a driver of the evolution of antibiotic resistance but not the mechanism of death in ampicillin-treated cells. (A) Levels of transcription of antioxidative genes in the wild type and the *ΔrecA* strain after the single treatment with ampicillin at 50 μg/ml for 8 hours. (B) Addition of 50 mM antioxidative compound GSH prevented the evolution of antibiotic resistance in the *ΔrecA* strain. (C) Survival fraction after the exposure to ampicillin at 50 μg/ml for 8 hours in the wild type and the *ΔrecA* strain with or without the addition of GSH at 50 mM. (D) Whole genome sequencing confirms the DNA mutations in the *ΔrecA* strain with or without the addition of GSH at 50 mM after the single treatment of ampicillin at 50 μg/ml for 8 hours.

Next, by comparing the expression of genes in biological processes of DNA replication, recombination and repair, SOS response, ABC transport system, and quorum-sensing system (Fig. 3A and Fig. S3), we found a series of genes related to the DNA damage response were slightly changed, but the transcription of gene *pinR* was considerably upregulated in the *ΔrecA* strain after the single treatment of ampicillin (Fig. 3A). The gene *pinR* encodes a putative site-specific recombinase PinR, which is a lambdoid prophage and exists in many bacterium types, including the *E. coli, Tetragenococcus halophilus, Lactococcus lactis, Salmonella enterica and Streptococcus* (16-19). Although the function of PinR is yet to be clear, it is predicated that PinR shares a similar structure with the DNA invertase (Fig. S4) (20,21). Recent studies showed that PinR an catalyse an inversion of a 177-bp DNA fragment in the *Streptococcus* acting as a bacterial recombinase (17). We asked whether the induction of prophage PinR facilitated the evolution of antibiotic resistance in the *ΔrecA* strain. To test it, we constructed a *recA/pinR* double deletion strain (*ΔrecA/pinR*) and treated them with ampicillin at 50 µg/mL for 8 hours. Interestingly, the evolution of antibiotic resistance was inhibited in the *ΔrecA/pinR* strain (Fig. 3B). We further explored a cyclic adaptive laboratory evolution (ALE) experiment in the *ΔrecA/pinR* strain, and found that the long-term treatment with ampicillin for 3 weeks no longer induced the evolution of antibiotic resistance in the *ΔrecA/pinR* strain, even though the establishment of antibiotic resistance was indeed determined in the wild type strain (Fig. 3C), which suggested that targeting PinR offered a way out of the dilemma whether to target RecA to suppress the SOS response towards preventing the antibiotic resistance.

**Figure 3.**
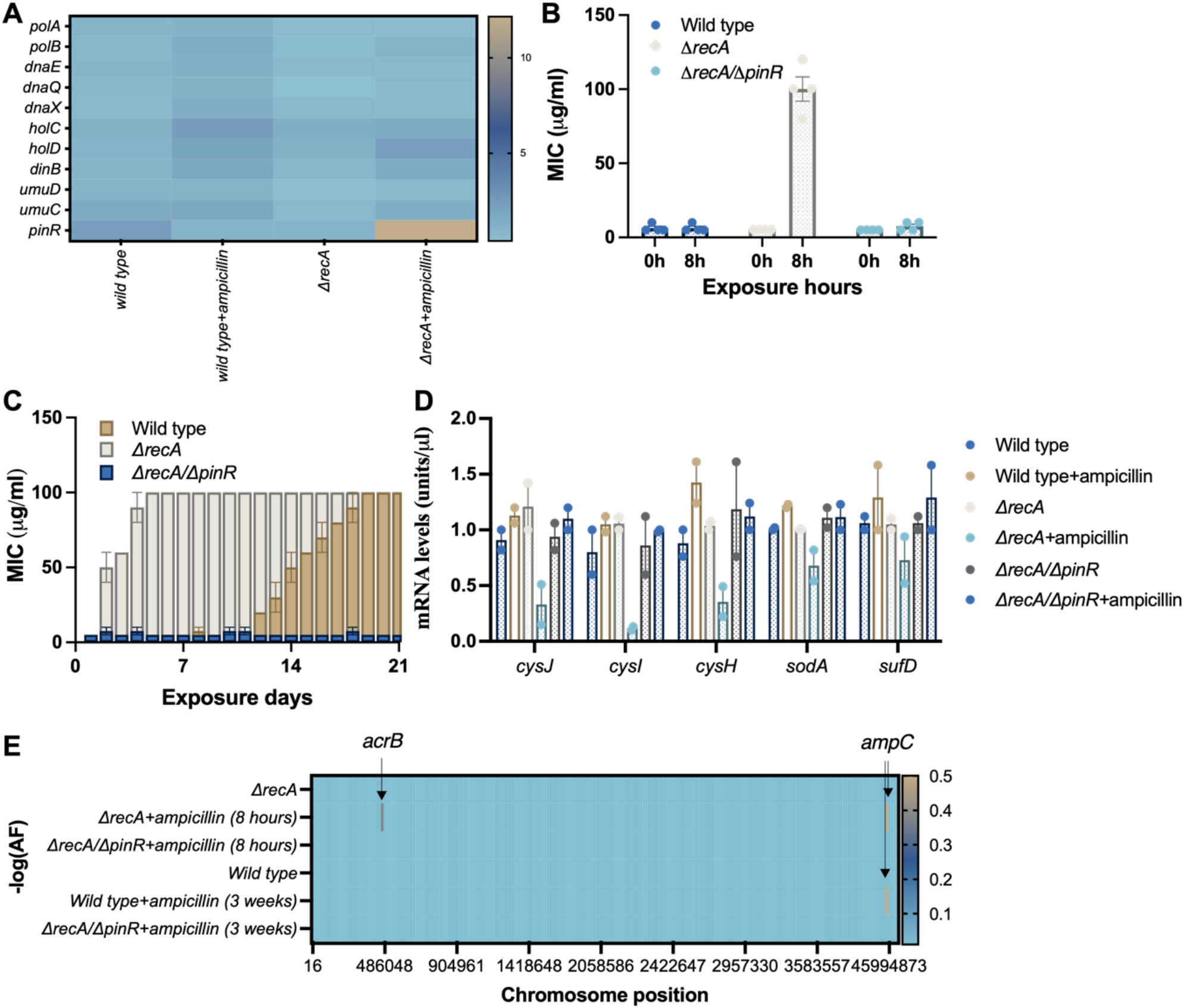
Induction of bacteriophage PinR facilitates the evolution of antibiotic resistance. (A) Levels of transcription of DNA replication, repair and recombination genes in the wild type and the *ΔrecA* strain after the single treatment with ampicillin at 50 μg/ml for 8 hours. (B) Deletion of *pinR* prevented the evolution of antibiotic resistance in the *ΔrecA/ΔpinR* strain after the single exposure to ampicillin at 50 μg/ml for 8 hours. (C) Deletion of *pinR* stoped the evolution of antibiotic resistance following an intermittent treatment with ampicillin at 50 μg/ml for 3 weeks in the *ΔrecA/ΔpinR* strain. (D) The mRNA levels of antioxidative genes in the in the wild type, *ΔrecA* and *ΔrecA/ΔpinR* strain after the single treatment of ampicillin at 50 μg/ml for 8 hours. (E) Whole genome sequencing confirms the DNA mutations in the *ΔrecA* and *ΔrecA/ΔpinR* after a single exposure to ampicillin at 50 μg/ml for 8 hours, or an intermittent treatment with ampicillin at 50 μg/ml for 3 weeks.

We further compared the mRNA levels of antioxidative genes including *cysJ, cysI, cysH, soda* and *sufD* by using ddPCR in the *ΔrecA*/*ΔpinR* strain and found that transcriptions of these genes were not affected by the single exposure to ampicillin (Fig. 3D). We finally sequenced the *ΔrecA*/*ΔpinR* surviving isolates and confirmed that the drug resistance-related DNA mutations were not detected in the *ΔrecA*/*ΔpinR* strain after a single or long-term treatment of ampicillin (Fig. 3E and Table S2). Collectively, these results indicated that the induction of prophage PinR played a role of “evolutionary potentiator” in facilitating the evolution of antibiotic resistance.

Although the role of ROS in the evolution of antibiotic resistance has been identified, our findings for the first time show that the induction of bacteriophage PinR can facilitate the evolution of antibiotic resistance in DNA repair deficiency bacteria through the suppression on the transcription of antioxidative genes and the thereafter ROS burst in cells (Fig. 4). These findings highlight the function of bacteriophage PinR that orchestrates the mutagenesis induced by the overaccumulation of ROS. Meanwhile, the SOS response is responsible for the induction of many lambdoid lysogens (22), thus the spontaneous SOS induction is proposed to trigger the induction of prophages (23). In addition, Little and Michalowski found that the intrinsic switching rate of *E. coli* lambda lysogens is almost undetectably low (<10^−8^/generation) in a *recA* mutant background indicating the spontaneous prophages induction has coevolved with specific triggers, *e*.*g*., SOS response and activation of RecA (24). However, to our knowledge, these results first show a different mechanism by which the antibiotic explore can induce the reactivation of prophages in DNA repair deficiency cells in a RecA-independent manner.

**Figure 4.**
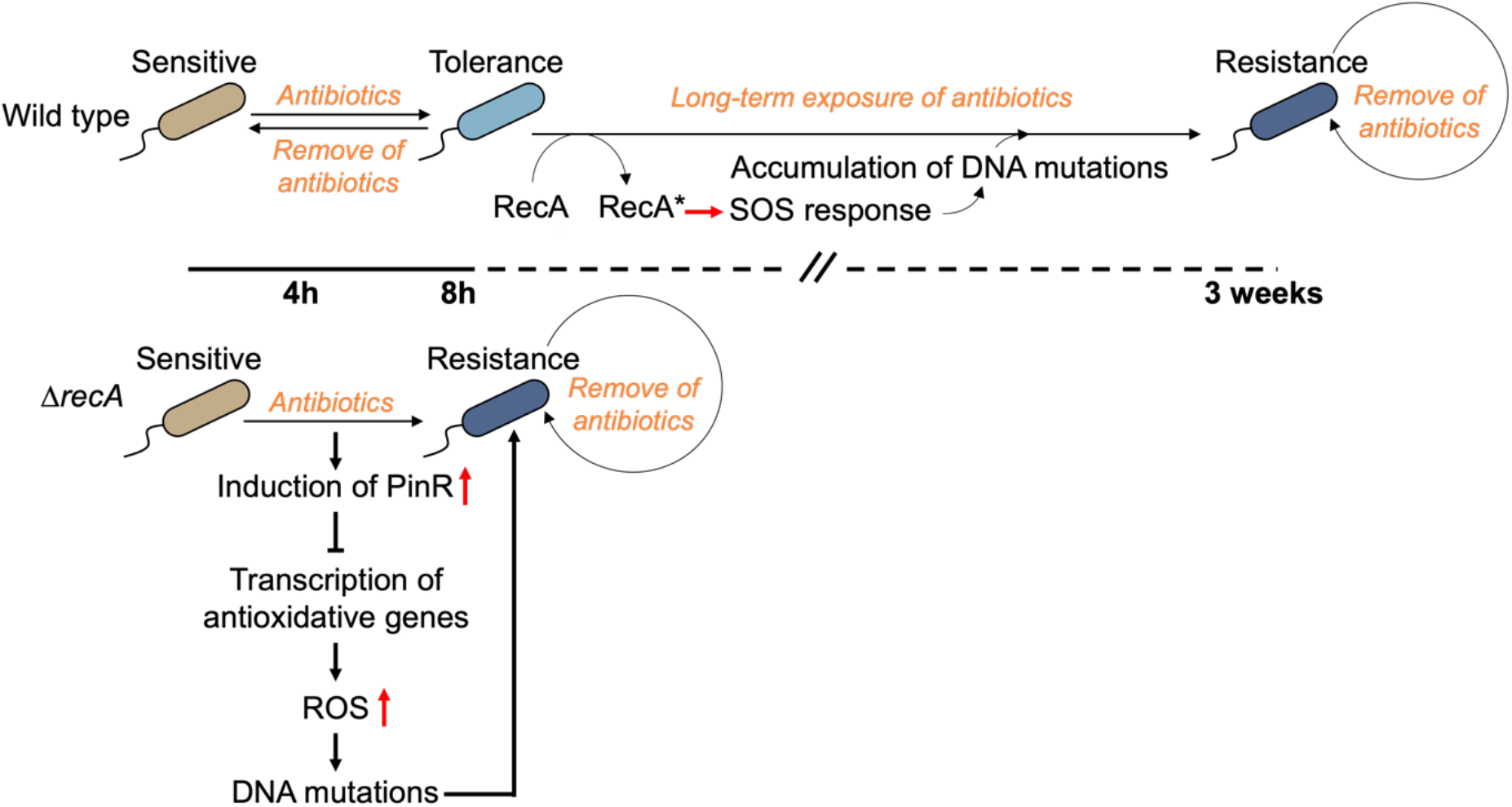
Mechanism of superfast evolution of antibiotic resistance. In the wild type strain, short-term exposure to antibiotics can induce the emergence of tolerance, and long-term treatment of drugs is able to trigger the bacterial evolution of antibiotic resistance from the tolerance. In this process, the induction of SOS response and the activation of its master regulator RecA contribute to the accumulation of drug resistance-related mutagenesis in cells. However, we here for the first time report that, in the *ΔrecA* strain, short-term exposure to the antibiotic can make bacteria evolve to be drug resistance in 8 hours. The induction of bacteriophage PinR induced by the single treatment with antibiotics plays a role of “evolutionary potentiator” in promoting the DNA mutations and facilitating the evolution of antibiotic resistance in the DNA repair deficiency bacteria through the repression on the transcription of antioxidative genes and the thereafter ROS burst in cells.

From a drug development perspective, strategies to directly target the transcription of “evolutionary potentiators” can combat the antibiotic resistance. Our results first demonstrate that deletion of *pinR* can avoid the rapid evolution of antibiotic resistance induced by the antibiotic exposure in DNA repair deficiency cells, while strategies to target RecA, *e*.*g*., the inactivation on RecA, can be safely implemented to disarm the bacterial resistance to other antibiotics. Moreover, with the role of PinR being clear, our evolutionary model can be generic, as the overproduction of ROS is a common action upon the treatment of many antibiotics and maintaining the transcription of oxidative stress response genes by inhibiting the “evolutionary potentiators” becomes a perspective approach to stop the stress-induced mutagenesis that is otherwise unavoidably induced by the treatments of other varieties of antibiotics. Therefore, our work suggests future studies on the “evolutionary potentiators” towards a safe and more effective strategy to be developed for infectious disease treatment.

## Acknowledgements

This work was supported by the Australia Research Council (ARC grant no: APP1165135, FT160100010), Science and Technology Innovation Commission of Shenzhen (KQTD20170810110913065), Australia China Science and Research Fund Joint Research Centre for Point-of-Care Testing (ACSRF658277, SQ2017YFGH001190).

## Author contributions

L.Z., G.P., Q.S., and D.J. designed the experiments. L.Z., Y.L., I.D., and D.J. wrote the manuscript with input from all co-authors. L.Z., G.P., and Y.L. conducted experiments and analysed data.

## Materials and Methods

### Bacterial strains, medium and antibiotics

Bacterial strains and plasmids used in this work are described in Table S3 and Table S4. Luria-Bertani (LB) was used as broth or in agar plates. *E. coli* cells were grown on LB agar (1.5% w/v) plates at 37°C, unless stated otherwise, antibiotics were supplemented, where appropriate. Whenever possible, antibiotic stock solutions were prepared fresh before the use.

### Treatment with antibiotics to induce evolutionary resistance

For the single exposure to antibiotic experiment, an overnight culture (0.6 mL; 1 × 10^9^ CFU/mL cells) was diluted 1:50 into 30 mL LB medium supplemented with antibiotics (50 μg/mL ampicillin, 1 mg/mL penicillin G, or 200 μg/mL carbenicillin) and incubated at 37°C with shaking at 250 rpm for 8 hours. After the treatment, the antibiotic-containing medium was removed by washing twice (20 min centrifugation at 1500 g) in fresh LB medium.

To test the capacity for tolerance, surviving isolates were immediately used or stored at -80°C for future use. To test resistance, the surviving isolates were first resuspended in 30 mL LB medium and grown overnight at 37°C with shaking at 250 rpm. The regrown culture was then plated onto LB agar supplemented with the appropriate selective antibiotics and incubated 16 hours at 37°C. Single colonies were isolated and used to test the resistance or stored at -80°C for future use.

For the intermittent antibiotic treatment experiments, an overnight culture (0.6 mL; 1 × 10^9^ CFU/mL cells) was diluted 1:50 into 30 mL LB medium supplemented with 50 μg/mL ampicillin and incubated at 37°C with shaking at 250 rpm for 4 hours. After treatment, the antibiotic-containing medium was removed by washing twice (20 min centrifugation at 1500 g) in fresh LB medium. The surviving isolates were resuspended in 30 mL LB medium and grown overnight at 37°C with shaking at 250 rpm. The killing treatment was applied as above to the regrown culture and repeated until resistance was established.

### Antibiotic susceptibility testing

The susceptibility of *E. coli* cells to antibiotics was measured by using minimum inhibitory concentration (MIC) testing (25). In brief, diluted overnight bacterial culture (10^7^ CFU/mL, determined by CFU counting) was used to inoculate wells of a sterile 96-well flat-bottomed plate. Various concentrations of antibiotics were added to the designated wells by serial dilutions with LB media to a final volume of 150 μL. Untreated controls were also included. The plate was incubated in the Synergy HT BioTek plate reader (BioTek Instruments Inc., USA) at 37°C with continuous moderate shaking to prevent biofilm formation (1800 rpm, amp. 0.549 mm x-axis) for 24 h and was programmed to measure the OD hourly at 595 nm (Gen5 software, BioTek Instruments Inc., USA). The MIC was defined as the lowest concentration of antimicrobial agent that inhibited 99% growth of *E. coli* when compared to the untreated control.

The capacity of tolerance was measured by using the minimum duration for killing 99% of the population (MDK_99_) testing (26). After the treatment with antibiotics, surviving cells were loaded in a 96 wells plate (approx. 10^4^ bacteria/well) that was filled with fresh LB medium supplemented with increasing amounts of ampicillin in which the lowest concentration of ampicillin was 100 μg/mL. Inoculation times are set so that all rows end their respective treatment at the same time. Once incubation is concluded, the plate is spun down to terminate the antibiotic exposure by washing away antibiotic remains and resuspending in fresh medium. The plate is then returned for overnight incubation. Empty wells indicate killing of > 99% of the population, because growth in the well would imply that at least one bacterium. An evaluation of MDK_99_ can be read directly from the plate depending on the treatment duration at which the plateau forms.

### Construction of deletion mutants

Lambda Red recombination was used to generate various gene deletions in *E. coli* strains followed by previous reported methods with modifications (27,28). Genomic DNA of *E. coli* K-12, containing insertions of a tetracycline or chloramphenicol resistance cassette to replace the open reading frame of *recA*, was used to make subsequent gene deletions. Primers (Table S5) were designed approximately 50 bp upstream and downstream to genes of interest on the chromosome, in order to amplify the tetracycline or chloramphenicol cassette as well as the flanking DNA sequence needed for homologous recombination. Phusion polymerase (NEB) was used to amplify DNA sequence (Table S5) and the reaction was cleaned up using a PureLink™ PCR purification kit (ThermoFisher Scientific) as per the manufacturer’s instructions. Background *E. coli* was made electro-competent and transformed with recombinase plasmid pKD46 and selected on LB agar plates containing 100 μg/mL ampicillin at 30°C. The strain now containing the plasmid was made electro-competent again using LB media containing 100 μg/mL ampicillin and 0.2% arabinose at 30°C. Amplified DNA was transformed into recipient strain by using 50 ng of DNA and 50μL of competent cells. Cells were allowed to recover in LB media for 1 hour at 30°C. Transformation was plated onto LB agar plates containing 10 μg/mL tetracycline or 17 μg/mL chloramphenicol and incubated overnight at 37°C. PCR was used to confirm insertion of the tetracycline or chloramphenicol resistance cassette at the correct site on the chromosome using primers upstream and downstream to the gene of interest. The newly constructed mutant strains were cured of plasmid pKD46 through incubation of LB streak plates at 42°C overnight. Loss of the plasmid was confirmed by lack of ampicillin sensitivity on LB agar plates. Mutant strains were made electro-competent and 50 μL of cells were transformed with plasmid pCP20 and incubated on 100 μg/mL ampicillin plates at 30°C overnight. A few colonies were then restreaked onto LB plates and incubated overnight at 42°C. Loss of cassette and plasmid was confirmed by PCR products.

### DNA extraction

Chromosomal DNA was extracted and purified using the PureLink™ Genomic DNA mini kit (ThermoFisher Scientific). In summary, a volume of 1 mL of overnight culture of the required strain was centrifuged for 2 minutes at 10,000 g. The cell pellet was resuspended in 180 μL PureLink™ genomic digestion buffer and 20 μL of Proteinase K and incubated at 55°C for 60 minutes until lysis was complete. A volume of 20 μL of RNase A (provided with the kit) was added to the lysate and mixed for 10 seconds on the vortex. The sample was then incubated for 2 minutes at room temperature. A volume of 200 μL of PureLink™ genomic Lysis/Binding buffer was added to the sample and mixed for 20 seconds by vertexing until a homogenous solution was obtained. 200 μL of 100% ethanol was added to the lysate and mixed well by vortexing for 5 seconds. The lysate (approximately 650 μL) was loaded onto a PureLink™ spin column in a collection tube and centrifuged at 10,000 g for 1 minute at room temperature. The flow through liquid was discarded and the column was washed twice with wash buffer 1 and wash buffer 2. The column was dried with a final spin for 3 minutes at 10,000 g. DNA was eluted using 200 μL of PureLink™ genomic elution buffer into a new 1.5 mL Eppendorf tube.

### Whole genome sequencing

The genomic sequencing was conducted following the Nextera Flex library preparation kit process (Illumina), and processed by Sangon Biotech, Shanghai, China. Briefly, genomic DNA was quantitatively assessed using Quant-iT picogreen dsDNA assay kit (Invitrogen, USA). The sample was normalised to the concentration of 1 ng/μL. 10 ng of DNA was used for library preparation. After tagmentation, the tagmented DNA was amplified using the facility’s custom designed i7 or i5 barcodes, with 12 cycles of PCR. The quality control for the samples was done by sequencing a pool of samples using MiSeq V2 nano kit - 300 cycles. After library amplification, 3 μL of each library was pooled into a library pool. The pool is then clean up using SPRI beads following the Nextera Flex clean up and size selection protocol. The pool was then sequenced using MiSeq V2 nano kit (Illumina, USA). Based on the sequencing data generated, the read count for each sample was used to identify the failed libraries (i.e., libraries with less than 100 reads). Moreover, based on the read count, libraries were pooled at a different amount to ensure equal representation in the final pool. The final pool was sequenced on Illumina NovaSeq 6000 Xp S4 lane, 2 × 150 bp.

### RNA extraction

RNA was extracted from the cell pellets using a PureLink RNA mini kit (Invitrogen) as per the manufacturer’s instructions. Briefly, 1×10^9^ log-phase bacterial cells were harvested and transferred to a microcentrifuge tube to centrifuge at 4°C for 5 minutes (500 g) to pellet cells. 100 μl of prepared lysozyme solution was added to the cell pellet to resuspend the cells by vertexing. 0.5 μl 10% SDS was then followed to be added and the cells were incubated at room temperature for 5 minutes. After the incubation, 350 μl lysis buffer prepared with 2-m-mercaptoethanol was added and cells were vortexed to mix well. Lysate was transferred to a 1.5 ml RNase-free tube and passed 5 times through a needle attached to an RNase-free syringe. The supernatant was collected through the centrifuge at 12,000 g for 2 minutes at room temperature. 250 μl 100% ethanol was added to each volume of bacterial homogenate and mixed thoroughly by vertexing to disperse any visible precipitate. Sample was then transferred to a Spin Cartridge and centrifuged at 12,000 g for 15 seconds at room temperature. Flow-through was discarded. 700 μl wash buffer I was added to the Spin Cartridge and centrifuged at 12,000 g for 15 seconds at room temperature. Flow-through was discarded. Wash buffer II was then added with ethanol to the Spin Cartridge and centrifuged at 12,000 g for 15 seconds at room temperature. Flow-through was discarded. After the washing, the Spin Cartridge was centrifuged at 12,000 g for 1 minute to dry the RNA attached onto the membrane to a Collection tube. 50 μl RNase-free water was followed to be added to the collection tube and all tubes were incubated at room temperature for 1 minute. All samples were centrifuged at 12,000 g for 2 minutes to collect the RNA, which was stored at -80°C for further use.

### Global transcriptome sequencing

The global transcriptome sequencing was processed and analysed by Genewiz, Jiangsu, China. Primers used in this work are listed in the Table S5.

### Droplet digital PCR (ddPCR)

Genomic DNA samples were added to the Bio-Rad 2 x ddPCR supermix at amounts of 0.05 ng DNA per 22 μL ddPCR reaction, according to the ddPCR Bio-Rad user manual. Samples were converted into droplets using a Bio-Rad QX200 droplet generator. After the droplet generation, the plate was transferred to a thermal cycler and reactions were run under the standard cycling conditions. After PCR, the plate was loaded onto the Bio-Rad QX200 Droplet Digital Reader, and data analysis was performed using Bio-Rad Quantasoft™ software. CNV analysis by ddPCR involves quantification of target and reference loci through the use of duplex target and reference assays. In QuantaSoftTM software, copy number is determined by calculating the ratio of the target DNA concentration to the reference DNA concentration, times the number of copies of reference species in the genome. The error bars on a CN estimate in QuantaSoftTM software are the 95% confidence interval of this measurement.

### Statistical analysis

Statistical analysis was performed using GraphPad Prism v.9.0.0. All data are presented as individual values and mean or mean ± s.e.m. A one-tailed unpaired Student’s t-test using a 95% confidence interval was used to evaluate the difference between two groups. For more than two groups, a one-way ANOVA was used. A probability value of *P* < 0.05 was considered significant. Statistical significance is indicated in each figure. All remaining experiments were repeated independently at least fourth with similar results.

---

**Figure S1.**
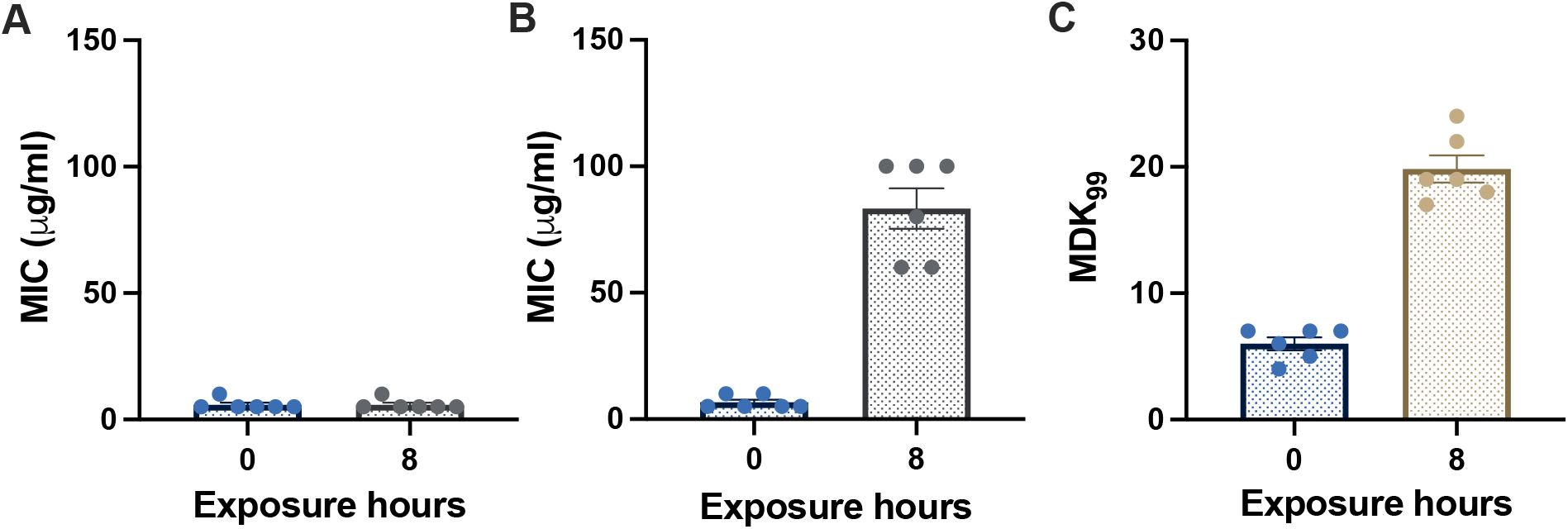
Superfast evolution of β-lactam resistance. The changes of MICs in the wild type (A) and the *ΔrecA E. coli* strain (B) after a single treatment with ampicillin at 50 µg/mL for 8 hours. (C) Changes of the MDK_99_ in wild type strain after the exposure to ampicillin at 50 µg/mL for 8 hours.

**Figure S2.**
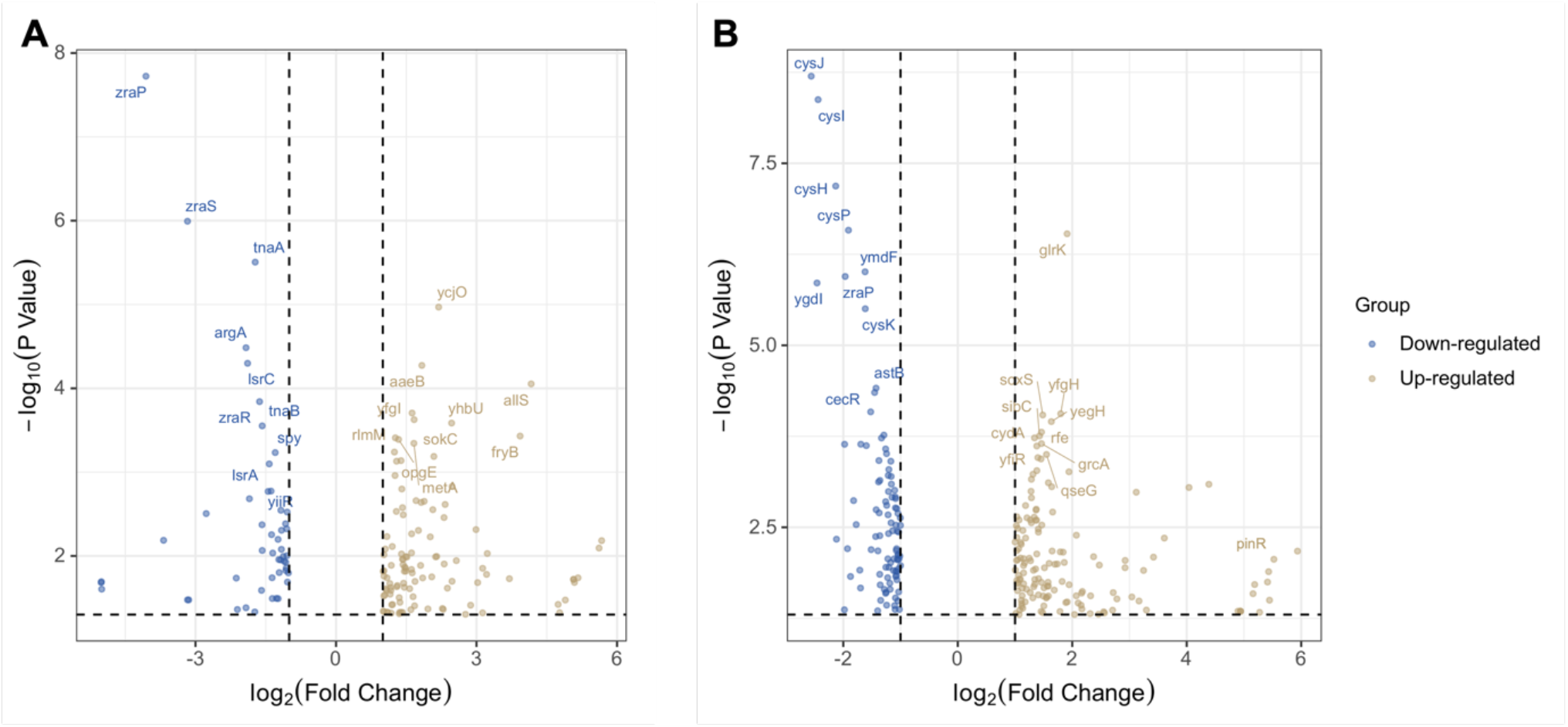
Summary of genome-wide expression changes in the wild type strain (A) and the *ΔrecA* strain (B) after the single exposure to ampicillin at 50 µg/mL for 8 hours. The top 10 most differentially expressed genes are labelled in each plot. Blue dots indicate genes with a significant downregulation compared to the untreated control (log_2_FC > 2 and *P* value < 0.05), and yellow dots indicate genes with a significant upregulation compared to the untreated control (log_2_FC > 2 and *P* value < 0.05).

**Figure S3.**
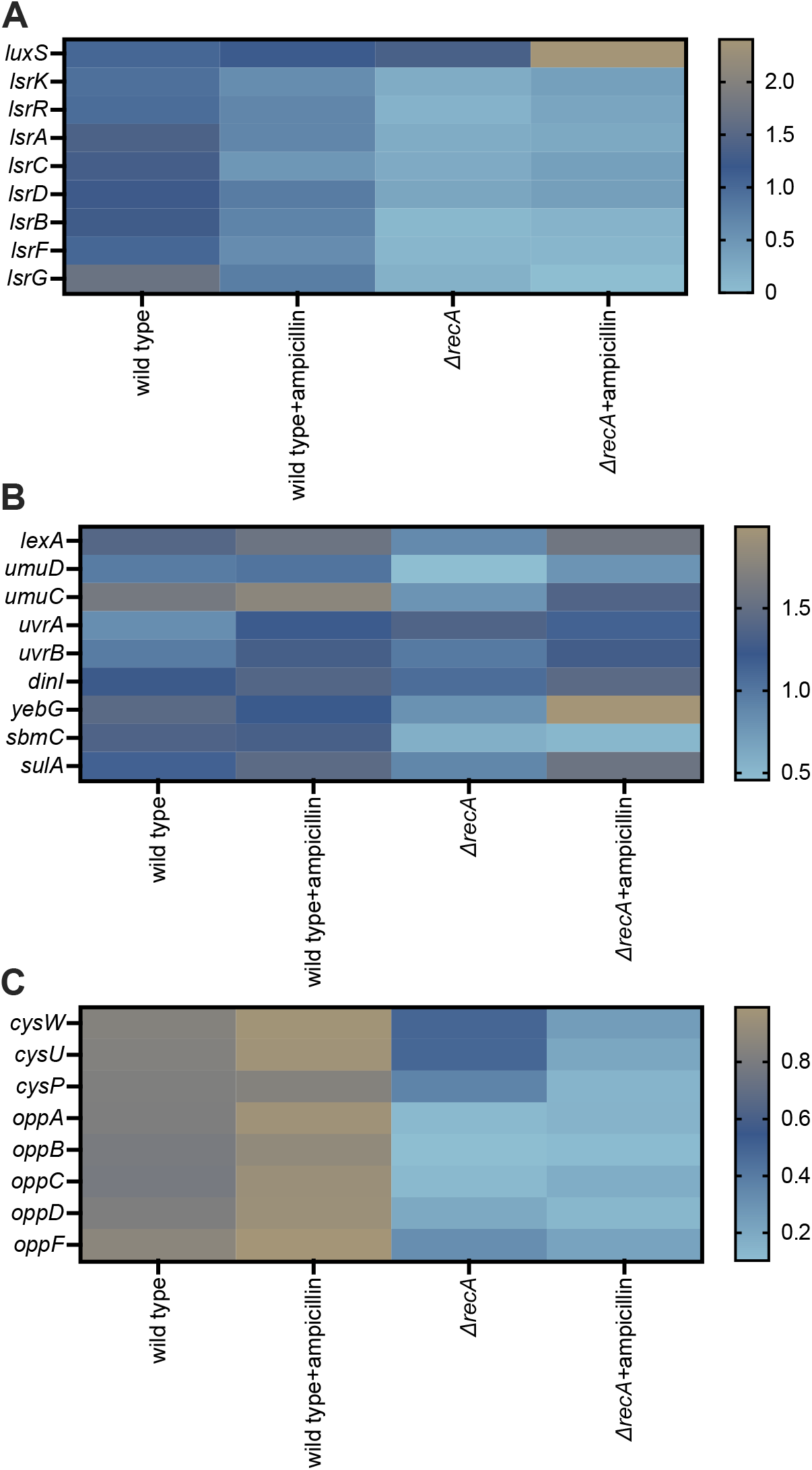
Heatmaps show log_2_FC data for each strain treated with or without single exposure to ampicillin at 50 μg/ml for 8 hours. (A) Quorum-sensing system genes. (B) SOS response genes. (C) ABC transport system genes.

**Figure S4.**
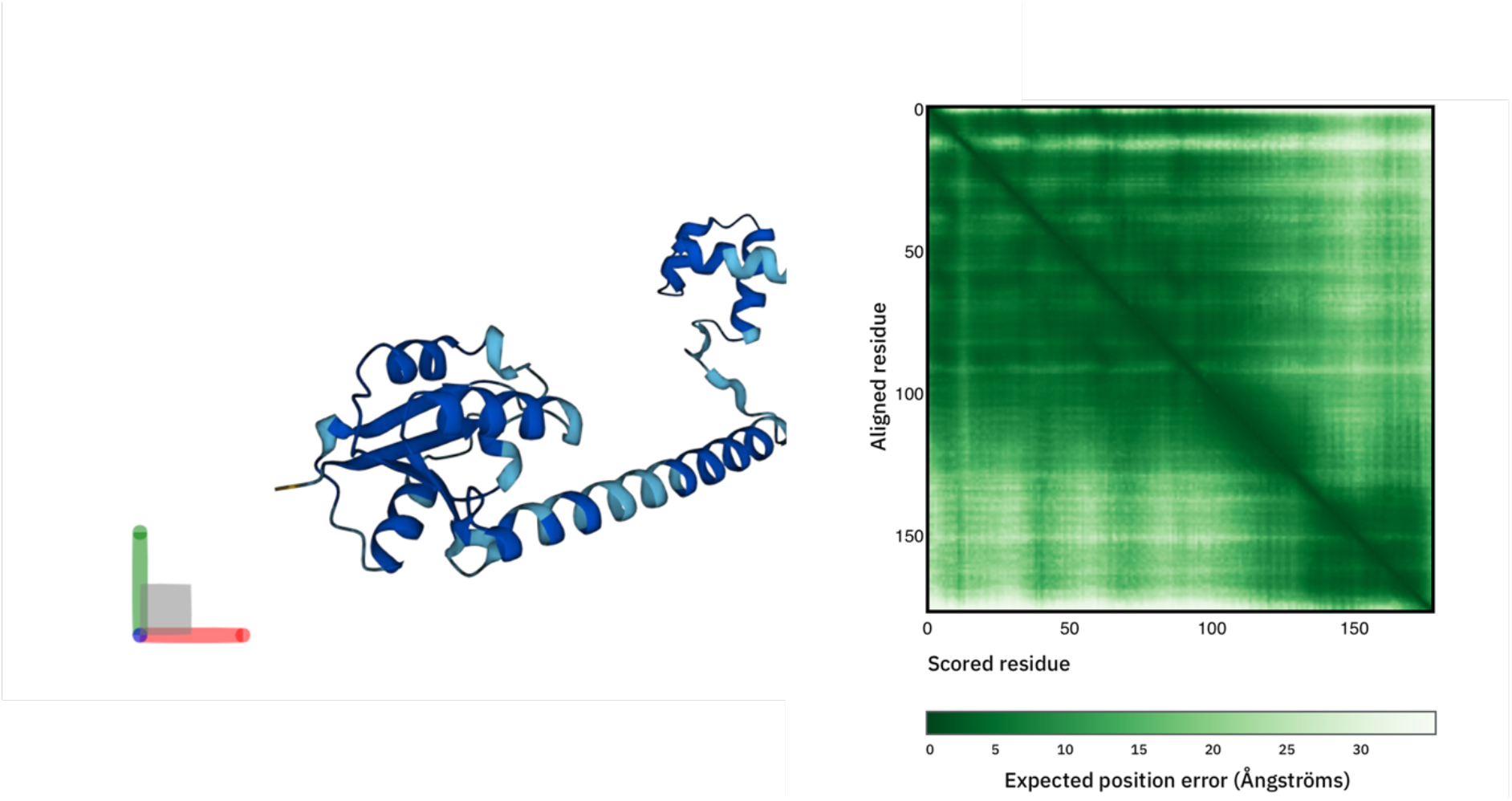
The proposed structure of DNA invertase PinR predicted by AlphaFold.

**Table S1.**
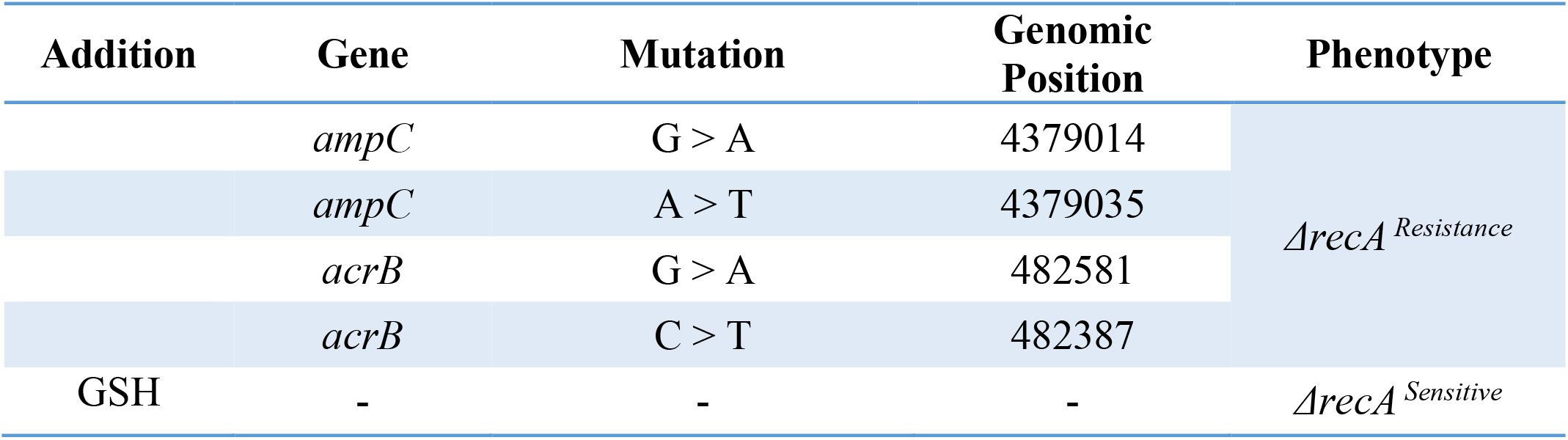
DNA mutations detected in the *ΔrecA* resistant isolates.

**Table S2.** DNA mutations detected in the wild type and *ΔrecA* resistant isolates.

| Exposure time | Gene | Mutation | Genomic Position | Phenotype |
| --- | --- | --- | --- | --- |
| 8 hours | *ampC* | G > A | 4379014 | *ΔrecA*<sup>Resistance</sup> |
| 8 hours | *ampC* | A > T | 4379035 |  |
| 8 hours | *acrB* | G > A | 482581 |  |
| 8 hours | *acrB* | C > T | 482387 |  |
| 3 weeks | *ampC* | G > A | 4379014 | Wild type<sup>Resistance</sup> |
| 3 weeks | *ampC* | + TA | 4379017 |  |
| 3 weeks | *ampC* | G > A | 4379014 |  |
| - | - | - | - | *ΔrecA/pinR*<sup>Sensitive</sup> |

**Table S3.** Strains used in this study.

| Strain | Relevant Genotype | Parent strain | Source |
| --- | --- | --- | --- |
| DH5α | - | - | Lab stock |
| *E. coli* K-12 | *recA*<sup>+</sup> *lexA*<sup>+</sup> | - | Lab stock |
| LZ101 | *lexA*<sup>+</sup> *ΔrecA::Tet* | K-12 | Lab stock |
| LZ102 | *ΔrecA::Tet*, *ΔpinR::Kan* | LZ101 | This study |

**Table S4.** Plasmids used in this study.

| Plasmid | Relevant Genotype | Parent strain |
| --- | --- | --- |
| pKD46 | Helper plasmid for Lambda RED recombination | Lab stock |

**Table S5.**
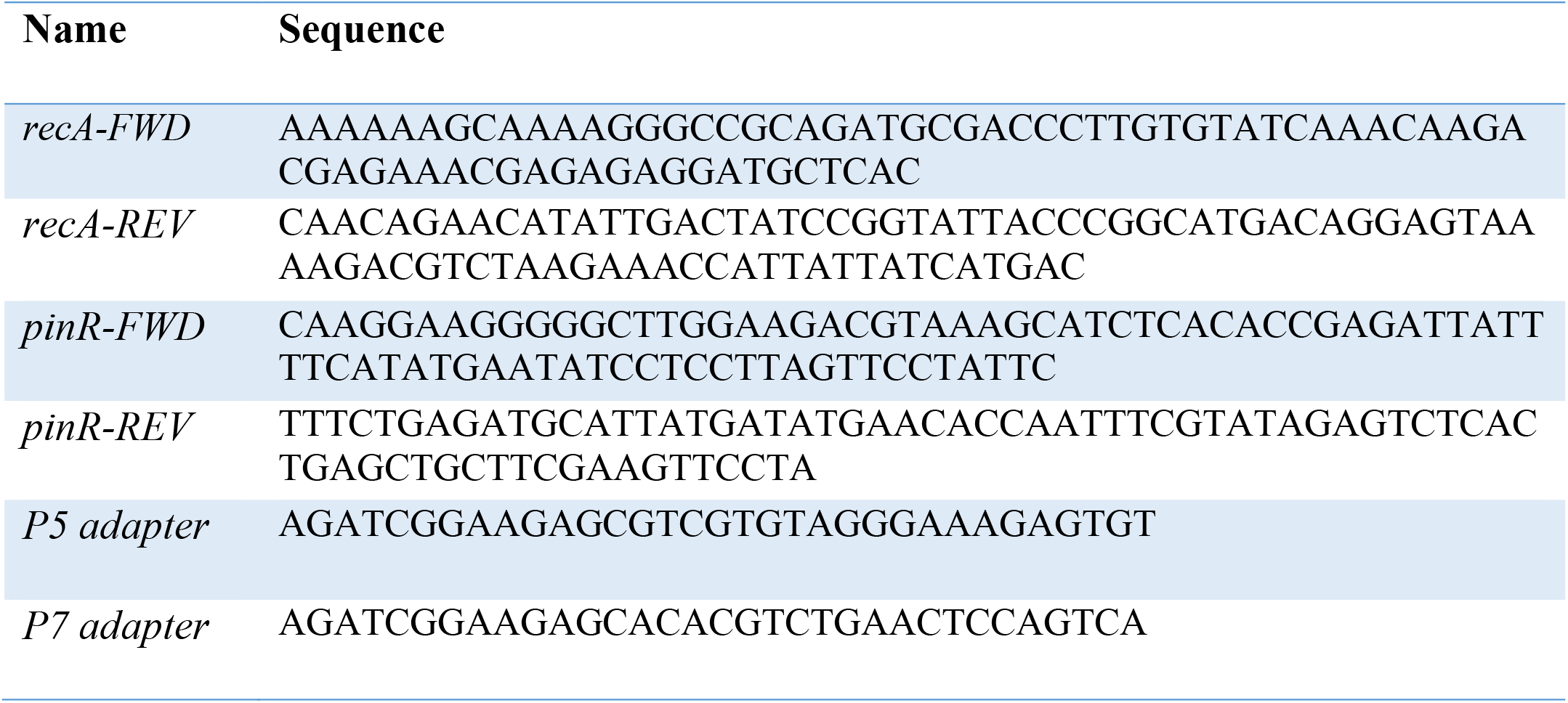
Primers used in this study.

## References

1. Rebecca S. Shapiro. Antimicrobial-Induced DNA Damage and Genomic Instability in Microbial Pathogens. PLOS Pathogens. 11, e1004678 (2015).

2. Darja Žgur-Bertok. DNA Damage Repair and Bacterial Pathogens. PLOS Pathogens. 9 PLOS Pathogens, e1003711 (2013).

3. Dale B. Wigley. Bacterial DNA Repair: Recent Insights into The Mechanism of RecBCD, AddAB and AdnAB. Nature Reviews Microbiology. 11, 9–13 (2013).

4. Alexander Harms, Etienne Maisonneuve, Kenn Gerdes. Mechanisms of Bacterial Persistence during Stress and Antibiotic Exposure. Science. 354, aaf4268 (2016).

5. Katarzyna H. Maslowska, Karolina Makiela-Dzbenska, Iwona J. Fijalkowska. The SOS System: A Complex and Tightly Regulated Response to DNA Damage. Environ Mol Mutagen. 60, 368–384 (2019).

6. Jason C. Bell, Stephen C. Kowalczykowski. RecA: Regulation and Mechanism of a Molecular Search Engine. Trends in Biochemical Sciences. 41, 491–507 (2016).

7. Michael M. Cox. Motoring Along with the Bacterial RecA Protein. Nature Reviews Molecular Cell Biology. 8, 127–138 (2007).

8. Michael M. Cox. The Bacterial RecA Protein as a Motor Protein. Annu. Rev. Microbiol. 57, 551–577 (2003).

9. Jiafeng Liu, Orit Gefen, Irine Ronin, Maskit Bar-Meir, Nathalie Q Balaban. Effect of tolerance on the evolution of antibiotic resistance under drug combinations. Science. 367, 200–204 (2020).

10. Irit Levin-Reisman et al. Antibiotic Tolerance Facilitates the Evolution of Resistance. Science. 355, 826–830 (2017).

11. Le Zhang et al. Superfast Evolution of Antibioitc Resistance in Escherichia coli. (2022). BioRxiv, doi: http://doi.org/10.1101/2022.03.29.486198.

12. Alexander Harms, Etienne Maisonneuve, Kenn Gerdes. Mechanisms of Bacterial Persistence during Stress and Antibiotic Exposure. Science. 354, aaf4268 (2016).

13. Sophie Helaine, Elisabeth Kugelberg. Bacterial Persisters: Formation, Eradication, and Experimental Systems. Trends in Microbiology. 22, 417–424 (2014).

14. Amy L. Spoering, Kim Lewis. Bio¢Lms and Planktonic Cells of Pseudomonas Aeruginosa Have Similar Resistance to Killing by Antimicrobials. J. Bacteriol. 183, 6746–6751 (2001).

15. Michael A Kohanski, Mark A DePristo, James J Collins. Sublethal antibiotic treatment leads to multidrug resistance via radical-induced mutagenesis. Mol. Cell. 37, 311–320 (2010).

16. Butland G, Peregrín-Alvarez JM, Li J, Yang W, et al. Interaction network containing conserved and essential protein complexes in Escherichia coli. Nature. 433, 7025:531-537 (2005).

17. Steward KF, Harrison T, Robinson C, Slater J, et al. PinR mediates the generation of reversible population diversity in Streptococcus zooepidemicus. Microbiology. 161, 5:1105-1112 (2015).

18. Johansson MHK, Bortolaia V, Tansirichaiya S, et al. Detection of mobile genetic elements associated with antibiotic resistance in Salmonella enterica using a newly developed web tool: MobileElementFinder. J Antimicrob Chemother. 76, 1:101-109 (2021).

19. Abriouel H, Pérez Montoro B, de la Fuente Ordoñez JJ, et al. New insights into the role of plasmids from probiotic Lactobacillus pentosus MP-10 in Aloreña table olive brine fermentation. Sci Rep. 9, 1:10938 (2019).

20. Jumper, J et al. Highly accurate protein structure prediction with AlphaFold. Nature. (2021).

21. Varadi, M et al. AlphaFold Protein Structure Database: massively expanding the structural coverage of protein-sequence space with high-accuracy models. Nucleic Acids Research (2021).

22. Nanda AM, Thormann K, Frunzke J. Impact of spontaneous prophage induction on the fitness of bacterial populations and host-microbe interactions. J Bacteriol. 197 (3):410–419 (2015).

23. Little JW. Chance phenotypic variation. Trends Biochem Sci. 15 (4):138 (1990).

24. Little JW, Michalowski CB. Stability and instability in the lysogenic state of phage lambda. J Bacteriol. 192 (22):6064–6076 (2010).

25. Eric M. Scholar, William B. Pratt. The Antimicrobial Drugs. Oxford University Press.

26. Asher Brauner, Noam Shoresh, Ofer Fridman, Nathalie Q. Balaban. An Experimental Framework for Quantifying Bacterial Tolerance. Biophysical Journal. 112, 2664–2671 (2017).

27. Kirill A. Datsenko, Barry L. Wanner. One-Step Inactivation of Chromosomal Genes in Escherichia Coli K-12 Using PCR Products. Proc Natl Acad Sci. 97, 6640–6645 (2000).

28. Tomoya Baba et al. Construction of Escherichia Coli K-12 In-Frame, Single-Gene Knockout Mutants: The Keio Collection. Mol Syst Biol, 2, 2006.0008 (2006).

